# Transient exposure of a buried phosphorylation site in an autoinhibited protein

**DOI:** 10.1101/2021.05.10.443419

**Authors:** Simone Orioli, Carl G. Henning Hansen, Kresten Lindorff-Larsen

## Abstract

Autoinhibition is a mechanism used to regulate protein function, often by making functional sites inaccessible through the interaction with a cis-acting inhibitory domain. Such autoinhibitory domains often display a substantial degree of structural disorder when unbound, and only become structured in the inhibited state. This conformational dynamics makes it difficult to study the structural origin of regulation, including effects of regulatory post-translational modifications. Here, we study the autoinhibition of the Dbl Homology domain in the protein Vav1 by the so-called acidic inhibitory domain. We use molecular simulations to study the process by which a mostly unstructured inhibitory domain folds upon binding and how transient exposure of a key buried tyrosine residue makes it accessible for phosphorylation. We show that the inhibitory domain, which forms a helix in the bound and inhibited stated, samples helical structures already before binding and that binding occurs via a molten-globule-like intermediate state. Together, our results shed light on key interactions that enable the inhibitory domain to sample a finely-tuned equilibrium between an inhibited and a kinase-accessible state.

## Introduction

Regulation of protein activity is crucial for cellular function, and the activity of a large number of proteins is regulated by *cis*-acting elements or modules (***Pufall and Graves, 2002***). When such modules interact with a protein in a way to inhibit its function, this is termed autoinhibition. This mechanism is widespread and can be employed to attenuate DNA-protein or protein-protein interactions (***Dombroski et al., 1993***; ***Ko and Prives, 1996***; ***Stöven et al., 2000***; ***Lee et al., 2005***), to regulate enzymatic activity (***Hubbard, 2004***) or even to impact protein localisation (***Pearson et al., 2000***). Schematically, autoinhibited proteins are usually composed of two main modules, an inhibitory module (IM) and a functional domain (FD), which is regulated by the IM itself (***Trudeau et al., 2013***). The inhibition can either be allosteric in nature or occur by physically hindering the access to the FD’s binding or catalytic site. As a consequence, autoinhibited proteins populate at least two distinct states: the inactive (autoinhibited) state, where the IM is bound to the FD, and an active state where the FD and IM do not interact directly.

Conformational transitions between the inactive and the active states of an autoinhibited protein often requires some degree of flexibility in the region that connects the IM with the FD (***Trudeau et al., 2013***). In some cases the IM itself has been reported to be intrinsically disordered in the active state, adopting a stable structure only upon binding to the FD (***Kim et al., 2000***; ***Pufall et al., 2005***), and it has been shown that autoinhibited proteins are enriched in intrinsic disorder (***Trudeau et al., 2013***). Among other important implications, this fact makes these systems useful examples to study so-called folding-upon-binding equilibria in intrinsically disordered proteins (IDPs) or regions (IDRs), a central challenge in structural biology (***Robustelli et al., 2020***; ***Malagrinò et al., 2020***).

IDPs and IDRs populate several structurally heterogeneous conformations when unbound (***Van Der Lee et al., 2014***; ***Habchi et al., 2014***), but are nonetheless often able to form highly specific interactions with their binding partners, often (but not always (***Schuler et al., 2020***)) through a process called folding-upon-binding (***Dyson and Wright, 2002***). The characterisation of the binding process at the atomic level would help provide insight into the process and a deeper understanding of how the physical and chemical properties of the IM and FD (sequence, charge distribution etc.) influence their biology. While for example NMR and single-molecule fluorescence spectroscopy has provided a wealth of details on the structural properties of bound and unbound states, and the kinetics of interconversion, detailed structural studies of the actual binding process are difficult to achieve by experiments (***Kim et al., 2018***; ***Sturzenegger et al., 2018***). On the other hand, all-atom molecular dynamics (MD) simulations have the ability to resolve binding processes at high spatial and temporal resolution, and have previously been used to characterize folding-upon-binding processes (***Piana et al., 2013***; ***Robustelli et al., 2020***; ***Henriques and Lindorff-Larsen, 2020***). In this context it is worth noting that the tethering of an ID to a FD may increase the local concentration of a disordered region (***Sgourakis et al., 2010***), thus making *cis*-acting regulatory regions particularly attractive targets to study such processes.

Here we focus on the Dbl Homology (DH) domain of the protein Vav1 as an example of a protein that is regulated via an intrinsically disordered IM. Vav proteins are large multi-domain proteins whose main function is to act as a guanine nucleotide exchange factor (GEF) for Rho family GTPases (***Bustelo, 2001***; ***Tybulewicz, 2005***). Vav proteins contain eight domains, of which the DH domain holds the GEF activity (***Abe et al., 1999***), with the remaining domains regulating function either through modulating activity or localization (***Bustelo, 2001***; ***Tybulewicz, 2005***). Just N-terminally to the DH domain is found the so-called acidic (Ac) domain, and the structure of Vav1 (***Aghazadeh et al., 2000***) revealed that part of the Ac domain blocks the active site in the DH domain forming an autoinhibited state (Fig. 1A). This autoinhibition is recapitulated in a minimal construct, termed AD, consisting of a short helical region of the Ac (the inhibitory module) and the DH (the functional domain) domains. This construct has a ∼10 fold decreased GEF activity compared to the DH domain alone (***Li et al., 2008***). The presence of additional domains in Vav decreases activity by an additional 10 fold, but NMR and biochemical experiments have shown that the AD construct captures the main mechanism of inhibition and activation, and that the remaining domains merely modulate these (***Yu et al., 2010***; ***Li et al., 2008***).

**Figure 1.**
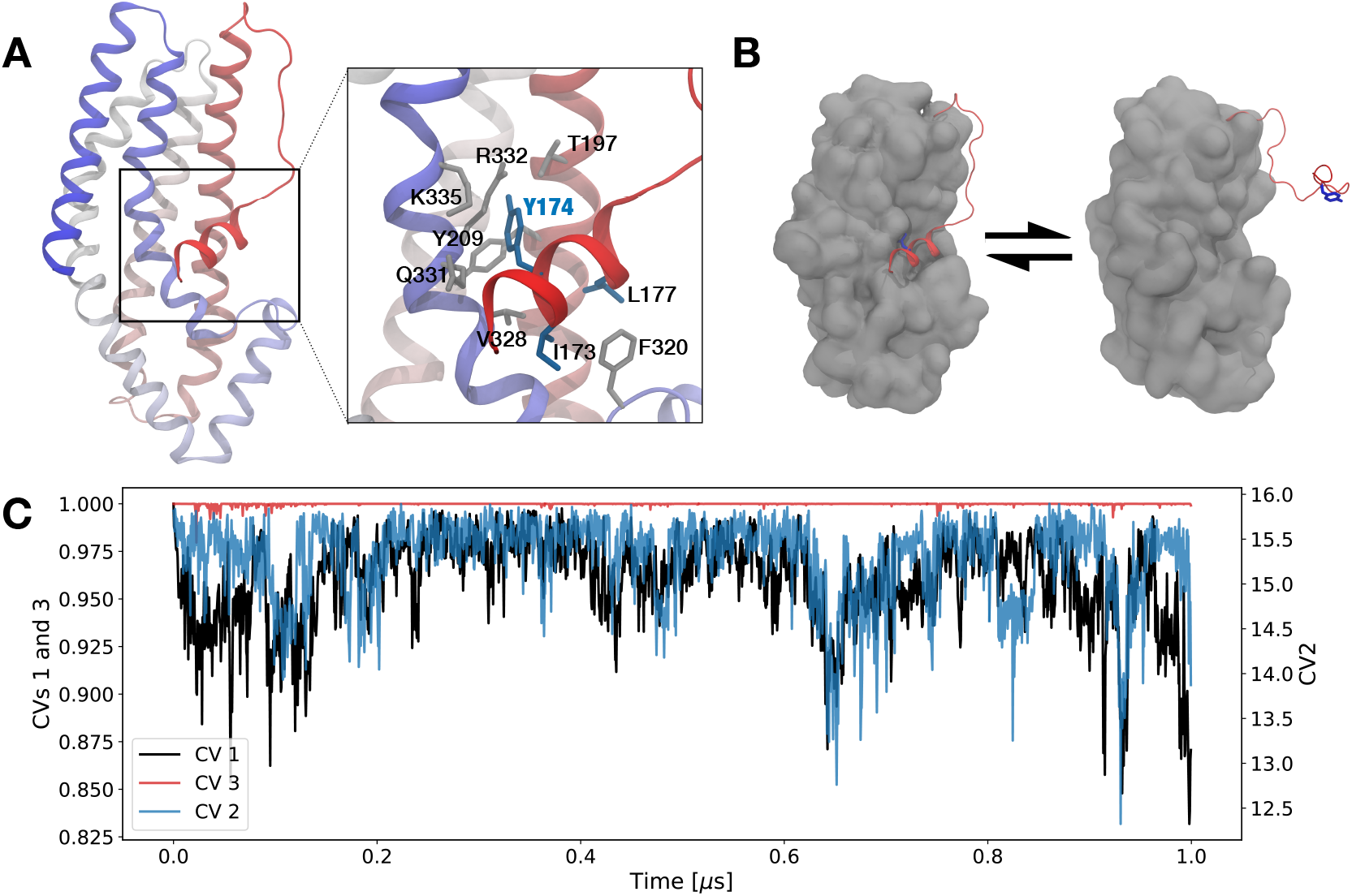
(A) Cartoon representation of the crystal structure of Vav1, coloured from the N-terminus (red) to C-terminus (blue). The box shows details with the short helix in the Ac domain, highlighting key residues in the interaction between the Ac and DH domains. (B) A schematic representation of the equilibrium between bound and unbound structures of the Ac inhibitory region. Y174 is depicted as blue sticks. (C) A time-series of the three collective variables (CV1–3) that we used to analyse the 1 *µ*s-long MD simulation started from the Vav1 crystal structure. CV1 quantifies the fraction of native contacts between the Ac-helix and the DH domain, CV2 probes the native contacts in the Ac-helix, and CV3 measures the fraction of native contacts between Y174 and the surrounding residues (within a 6Å radius in the crystal structure).

Activation of Vav1 occurs by phosphorylation of Y174 (Fig. 1A), located centrally in the Ac region. In the autoinhibited state, Y174 is however buried in the interface between the Ac and DH domains (Fig. 1B), and thus inaccessible to the Src-family kinase that phosphorylates it. Through a series of NMR and biochemical experiments, it was previously shown that the Ac region—including Y174— exists in a dynamic equilibrium between a buried state and an exposed, kinase accessible state with a 10:1 population ratio (***Li et al., 2008***). When Y174 is phosphorylated, the active state becomes dominant, thus explaining the 10-fold increase in activity upon phosphorylation. These observations strongly suggest a regulatory mechanism in which the transient exposure of at least part of the Ac region leads to access of the kinase to Y174; once phosphorylated, the Ac domain remains dissociated and the protein is active (Fig. 1B). The dynamics of the Ac helix in the protein’s active state and the details of the conformational change, however, remain elusive and unexplained.

We have here examined the transient exposure of the autoinhibitory domain in Vav1 through MD simulations. In particular, we have asked the question whether it is possible to observe such slow and finely tuned motions through all-atom simulations with the goal to shed light on the mechanism behind them. The NMR experiments showed that the closed/open equilibrium is fast on the NMR timescale, yet sufficiently slow to give rise to substantial relaxation dispersion, suggesting that the process occurs in the microsecond to millisecond timescale (***Li et al., 2008***). Such slow processes are often important biologically, but remain difficult to study by simulations. It has previously been shown that enhanced sampling simulations can be used to capture slow motions on the microsecond to millisecond timescale. In particular, MD simulations coupled with metadynamics (***Bussi and Laio, 2020***) were able to capture motions observed by NMR in both Cyclophilin A (***Papaleo et al., 2014***) and the L99A variant of T4 lysozyme (***Wang et al., 2016***). In both cases, however, detailed structural information was available for both the dominant state as well as the transiently populated state, and this information was used to guide the simulations. Similarly to the recently studied ternary complex p27 (***Henriques and Lindorff-Larsen, 2020***), however, for Vav1 less is known about the structural origin underlying the dynamic motions, so no conformation is available for the active state of Vav1 that can be employed to simplify the sampling.

We collected over 60 *µ*s of sampling combining unbiased MD and metadynamics simulations to reveal how the binding of the inhibitory module to the DH domain occurs through a dominant pathway where the short AC-helix (residues 169-177) forms most of its internal contacts before binding to its hydrophobic binding pocket. This provides an indication that, in the active state, the inhibitory module of Vav1 lives in an equilibrium between a helical and an unfolded state and it is therefore able to retain secondary structure without binding to the DH domain. Moreover, we show through short MD simulations that the AD interface is dynamic on the microsecond timescale and the Ac-helix is in a fast equilibrium between a tightly bound and a molten globule-like pre-bound state.

## Results and Discussion

### The Ac helix is bound to the DH domain during microsecond MD simulations

We first performed multiple 1 *µ*s-long MD simulation of the Vav1 AD construct, consisting of the Ac region and the DH domain, and monitored the structure and dynamics of the two domains and the interface between them. We carried out two simulations (with two different force fields) starting from the crystal structure and two from the NMR structure. We observe substantial dynamics in simulations initiated from the NMR structure (Fig. S1, Fig. S2 and discussion in the supplementary material), in apparent disagreement with NMR experiments suggesting the inhibited state to be stable on this time scale. In contrast, simulations initiated from the crystal structure were stable, and we thus focus on those below.

Calculations of the root-mean-square fluctuations, the fraction of native contacts and visual inspection of the trajectory revealed the protein to be very stable with only substantial motion in a loop region and in the tip of the N-terminal region (Fig. S1). The Ac region and the AD interface were also overall very stable as monitored by three distinct collective variables (see Supporting Information for a detailed description): CV1 which quantifies the fraction of native contacts at the interface between the Ac and DH domains in AD, CV2 quantifies the amount of helical structure in the Ac-helix, and finally CV3 quantifies the interactions between the side chain in residue Y174 (whose phosphorylation leads to activation of Vav1) and residues in the binding site (T205, Y209, V328, Q331 and R332). As expected from the NMR experiments (***Li et al., 2008***), the behavior of these collective variables (especially CV3) confirms that Y174 remains firmly bound during the simulation (Fig. 1C).

### Metadynamics simulations show the AD complex is formed through a dominant binding pathway

To be able to probe the free energy landscape describing the motions involved in the transient exposure of Y174, we turned to enhanced sampling simulations. Indeed, the NMR experiments (***Li et al., 2008***) suggest that the motions of the inhibitory module occur on the *µ*s to ms scale: this goes beyond the timescales that can be easily explored using standard MD simulations. Based on previous experience studying processes on similar time scales (***Papaleo et al., 2014***; ***Wang et al., 2016***; ***Henriques and Lindorff-Larsen, 2020***) we opted for metadynamics as an enhanced sampling tool to accelerate the exploration of the Ac-helix binding-unbinding equilibrium. Metadynamics employs a time-dependent potential, defined along a set of predetermined CVs, to accelerate the sampling of rare events. The choice of CVs is critical and far from trivial, and here is further complicated by the fact that little information is available on the unbound state of the Ac helix. For our first metadynamics run, the natural choice was therefore to bias along CV1 and CV2, as a similar choice of CVs has been previously employed to analyse folding-upon-binding dynamics of IDPs onto folded domains (***Robustelli et al., 2020***). During this simulation (see Fig. S3A-C) we were able to observe multiple binding-unbinding events during the first 4 *µ*s, but the Ac-helix did not rebind correctly for the remaining 6 *µ*s of the run, and therefore the simulation failed to converge to a meaningful free energy landscape (see also Fig. S3D and the related discussion).

In order to enhance the sampling along slow degrees of freedom that were not included in this first metadynamics run, we used parallel bias metadynamics (PBMetaD, ***Pfaendtner and Bonomi (2015***)) to enable us to use more than two CVs. Indeed, it has been previously shown that this approach may be used to capture slow motions in proteins when many CVs need to be employed (***Heller et al., 2017***; ***Prakash et al., 2018***; ***Fu and Pfaendtner, 2018***; ***Henriques and Lindorff-Larsen, 2020***). In contrast to standard well-tempered metadynamics, where Gaussian hills are deposited on a multi-dimensional grid with dimension of the number of CVs used, PBMetaD works by introducing multiple one-dimensional and time-dependent biases, one per CV. In this way, sampling is enhanced but the computational efficiency of the simulation is not severely affected by the number of CVs used. At the same time, one dimensional free energies can be calculated from the sum of the deposited Gaussian hills, while the multidimensional free-energy surface can be determined via reweighting techniques (***Pfaendtner and Bonomi, 2015***). We employed five CVs (CV1–3 and two others, capturing different aspects of the Y174 binding and Ac-helix folding and unfolding dynamics; see the supplementary information for further details) and two parallel walkers in our PBMetaD run in order to enhance the sampling of unbinding and rebinding conformational transitions. We collected a total of 10 *µ*s of sampling (5 *µ*s per walker), during which we were able to sample multiple unbinding-rebinding events (Fig. S4 and S5). We note that although CV3 consists of a subset of the contacts contained in CV1, we hypothesize that explicitly biasing also this collective variable was important in aiding the sampling. In particular, the contacts formed by Y174 only account for a small portion of the contacts used in CV1, but still (as we show below) the specific orientation of Y174 is fundamental in determining whether the Ac-helix is properly bound. Thus, simultaneously using both CV1 and CV3 aids in defining both global and more local properties of the binding of the Ac-helix and, as shown below, contain independent information. We performed a block error analysis (***Flyvbjerg and Petersen, 1989***; ***Bussi and Tribello, 2019***) in order to assess the convergence of the CVs in the PBMetaD simulation, and this analysis shows that the estimated errors plateau at block sizes around 0.5 *µ*s (Fig. S6A) and indicate the convergence of the simulation. This is furthermore corroborated by the fact that we observe binding and unbinding transitions also towards the end of the simulations when only little bias is added (Fig. S6B–F).

Our PBMetaD simulations of the Vav1 AD construct resulted in a complex free energy surface with multiple minima and distinct states between fully unbound and bound structures (Fig. 2A). The main binding pathway, as indicated by the free energy landscape along CV1 and CV2, starts from an extended state where the the Ac-helix is unbound and unfolded (Fig. 2A, **1**) and progresses through a state with increased helical content in the 169-178 region (Fig. 2A, **2**). Binding of the Ac region to the DH domain is initiated by the insertion of I173 into the hydrophobic pocket formed by F320, L325 and V328 (Fig. 2A, **3** and inset), an interaction which stabilises the positioning of the helix close to the binding site. The formation of the fully bound AD complex progresses through the establishment of native contacts between Y174 and T205, Y209, V328, Q331 and R332 (Fig. 2B, **3** and **4**), forming a molten globule-like state with the helix locked in place (Fig. 2A, **4**). The final bound conformation (Fig. 2A, **5**) is reached through the consolidation of the remaining contacts, especially in the elbow region (M178, T205 and K208), while Y174 ultimately settles into the hydrophobic binding pocket (Fig. 2B, **5**).

**Figure 2.**
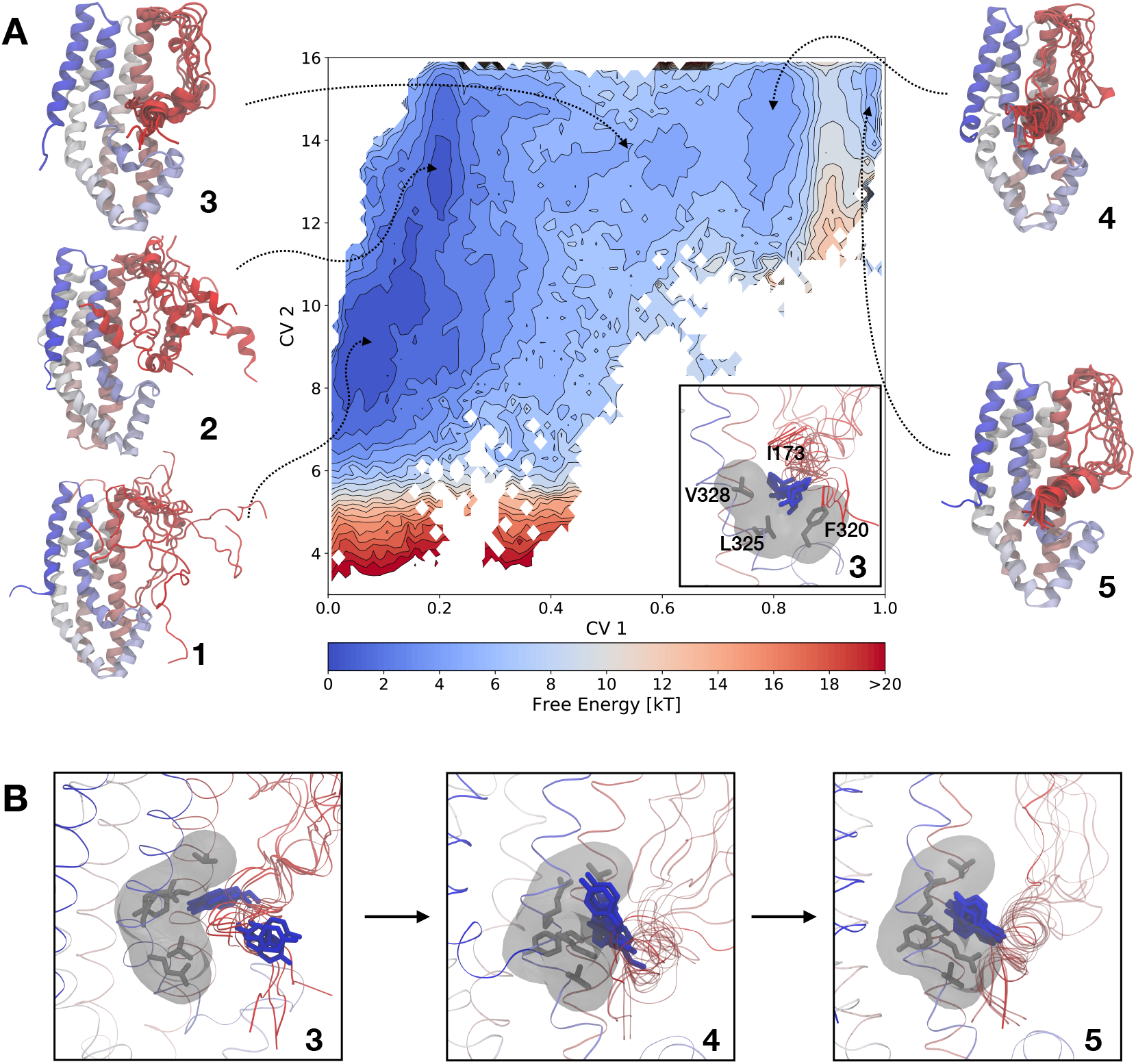
Free energy landscape and binding mechanism for the Ac helix to the DH domain. (A) Free energy surface, projected along CV1 and CV2, obtained from the dual-walker PBMetaD simulation. The minimum of the free energy surface has been set to zero energy, and low energy regions are indicated in blue, whereas high energy regions are shown in red. The ensembles (**1**–**5**) with 10 protein structures were extracted from the free energy region indicated by the dashed arrows (details in supporting information) and are used to help visualize the process of binding. The inset highlight residue I173 in representative configurations extracted from ensemble **3**. (B) Zoom in on residue Y174 (blue) for representative configurations extracted from ensembles **3**–**5**.

While examining the free energy landscape along CV1 and CV2 provides an intuitive overview of the binding process, the binding mechanism of the Ac inhibitory module is a complex process on a the high-dimensional surface. For this reason, it is important examine the process from the angle of multiple CVs. In the free energy landscape projected onto CV1/CV3 (Fig. 3A) and CV2/CV3 (Fig. 3B) we observe (i) that initial positioning of Y174 (CV3) can occur even with imperfect docking of the Ac-helix (CV1) (Fig. 3A) and (ii) that binding of Y174 (CV3) may occur with the Ac-region either helical or not (CV2) (Fig. 3B). This means that pathways exist (leftmost on the CV1/CV3 and CV2/CV3 landscapes) through which the AD complex can be rendered inactive by sequestering Y174 from the solvent, without the need for the inhibitory module to reach a fully bound conformation. As discussed above, while there is overlap between the contacts used in CV1 and CV3, they contain independent information. In particular, when the Ac-helix is (partially) unbound, CV1 and CV3 can vary independently (Fig. 3A) and we speculate that the inclusion of both CV1 and CV3 helped enhance sampling the binding events. Finally, for sake of completeness, we also report the one-dimensional free energies along the remaining two CVs used in the PBMetaD simulation (Fig. S7).

**Figure 3.**
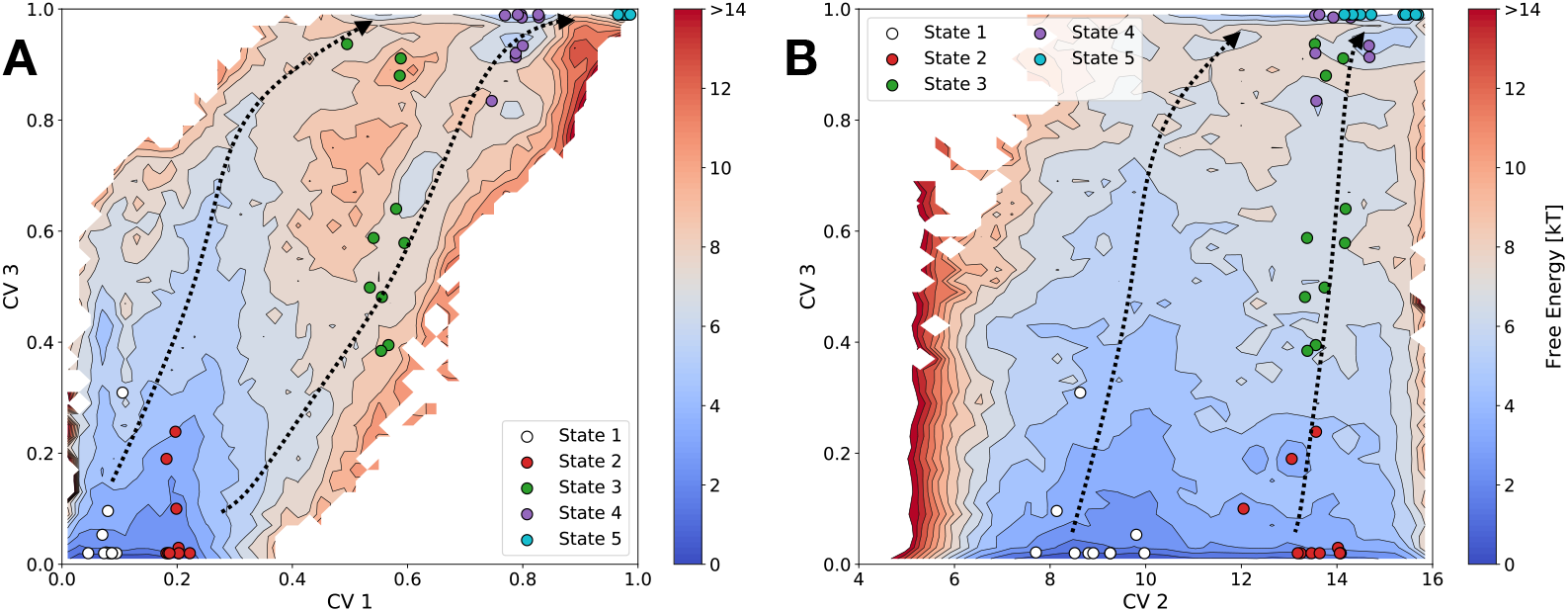
Free energy surface, projected along (A) CV1 and CV3 and (B) CV2 and CV3, obtained from the dual-walkers PBMetaD simulation. The minimum of the free energy surface has been set to zero energy, and low energy regions are indicated in blue, whereas high energy regions are shown in red. The coloured circles on the two plots indicate the positions of the representative configurations in Fig. 2 projected on to the corresponding free energy landscapes. The dashed arrows highlight the main binding pathways and are meant only as a guide for the eyes.

Summarising, visual inspection of the free energy landscape (Fig. 2A) suggests that the formation of the bound AD complex mainly progresses through a pathway where the Ac-helix consolidates most of its helical contacts while the the inhibitory module and the DH domain do not interact, and then binds to the hydrophobic patch formed by T205, Y209, F320, V328, Q331 and R332 (see supplementary video 1). Our simulations suggest that the inhibitory helix may fold independently of the interaction with the binding partner, i.e. the DH domain, and thus does not appear to follow an induced fit/folding-upon-binding mechanism. Indeed, such a mechanism would appear in the free energy landscape as either diagonal or L-shaped pathways in the bottom-right hand side of the CV1-CV2 plot. Moreover, the analysis of the free energy landscapes show that multiple binding pathways exist that make Y174 inaccessible to phosphorylation by Src kinase (Fig. 3).

### Short MD simulations confirm the binding pathway predicted by PBMetaD

As an enhanced sampling technique, the convergence of metadynamics simulations depends on the choice of CVs. If important CVs are not included one may still drive the transition via the bias potential, but simulations will not converge and transitions will not be observed as the bias is decreased. Presumably, this is what occurred in our initial set of simulations that used only CV1 and CV2 (see Fig. S3A-C). In contrast, our PBMetaD simulations appear to converge and we observe transition also towards the end of the simulations when the bias had decreased (Fig. S6). Nevertheless, to validate our PBMetaD simulations and the conformational states, we selected several configurations along the binding pathway and used them as a starting point for multiple unbiased MD simulations (each between 300 or 500 ns in length, see supporting information for further details). One of these, for example, showed a transition from a pre- to fully-bound structure (Fig. 4) while other sampled various regions of the conformational space of Vav1 (Fig. S8).

**Figure 4.**
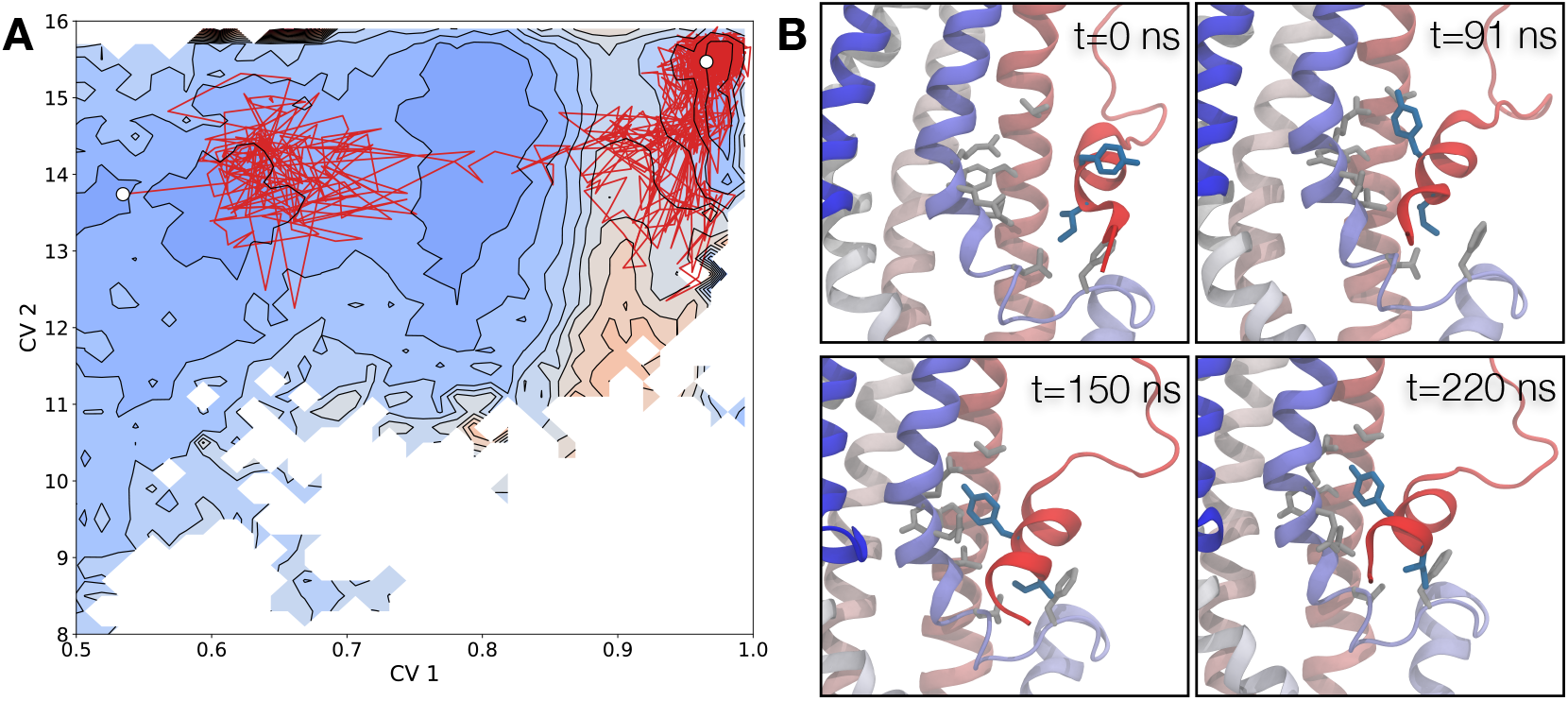
Binding of the Ac-helix in an unbiased molecular dynamics simulation. (A) Projection of an unbiased MD simulation, shown as a red solid curve, on to the free energy landscape from PBMetaD and defined by CV1 and CV2. (B) Zoom in on residue Y174 (blue) for selected frames along the binding trajectory.

We first attempted to analyse these simulations by building a Markov State Model (MSM) (***Chodera and Noé, 2014***) to obtain an overall view of the kinetic process of binding of the Ac-helix. We, however, failed to obtain a meaningful kinetic model because none of the simulations sampled the transition from state **2** to **3**, which evidently is a key limiting step in the overall binding process. We discuss our results for the MSM further in the supplementary material (Fig. 9) and, in absence of a reliable kinetic model, we proceeded to analyse the unbiased simulations as discussed below.

The behaviour of the short MD simulations, when projected onto the PBMetaD landscape along CV1 and CV2, strikingly agrees with our expectations, thus supporting the results from the PBMetaD simulation. In particular, they either sample the local and stable conformational states predicted by PBMetaD or follow low free energy pathways to sample multiple states (Fig. S8). These simulations also showed an unbinding event starting from a partially bound structure (Fig. S8D), and even the transition to a fully bound structure (Fig. 4A). Visual inspection of these simulations (supplementary videos 2 and 3), further support our interpretation of the binding pathway provided by PBMetaD. The sequence of events that result in the binding of the the Ac helix and Y174 to the binding pocket is as follows (see also Fig. 4B): At *t* = 0 ns, the helix is held close to the binding site by the interaction of I173 with the F320, L325 and V328 hydrophobic pocket, while Y174 is tilted pointing outwards with respect to the binding site; a rotamer jump around *t* = 90 ns brings Y174 closer to the native pocket and the corresponding interaction between Y174 and R332 is established shortly after (*t* = 150 ns). Finally, around *t* = 220 ns, the N-terminus of the helix tilts upwards to reach the native conformation. In addition to this structural view of the binding process, we conclude that the overall behaviour of MD simulations projected onto CV1 and CV2 is in agreement with the shape of the free energy profile obtained from PBMetaD, further validating that it represents the dynamics of Vav1.

Inspection of the free energy landscape from the PBMetaD simulations reveals a barrier (∼ 3.5kT, approximately symmetric on both sides) in the CV1-CV2 free energy surface, which separates the molten globule state **4** from the fully bound state, **5** (CV1 ∼ 0.95). We hypothesized that this would correspond to a transition state between state **4** and the fully bound structure **5**, and thus selected five conformations with CV1 ∼ 0.95 and CV2 values ranging from 14.3 to 15.4 and ran five independent unbiased MD simulations starting from each of these structures. We let each of these 25 trajectories run until they either reached state 5 (CV1 ≥ 0.98) or state 4 (CV1 ≤ 0.82) and evaluated the binding probability (*p*_binding_) as the ratio between the number of simulations reaching state **5** and the total number of trajectories. In good agreement with the PBMetaD predictions, we found *p*_binding_ = 0.48±0.05 (see supporting material for the estimation of the uncertainty), consistent with the value 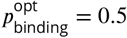 expected for a transition state (***Du et al., 1998***).

We also used the simulations initiated from the transition state region to estimate the barrier-crossing (transition path, TP) time and, based on this, the mean first passage (MFP) time between the two states. To this end, we considered only the reactive portions of the 25 simulations and estimated the forward, 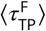, and backward, 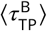, mean transition path times by averaging the time lengths of the, respectively, **4→5** and **5→4** transition paths. We found 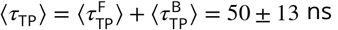. We then estimated the MFP time based on the relationship between the mean TP time and the MFP time: (***Hummer, 2004***; ***Chung et al ., 2009***)

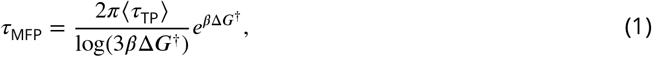

Here Δ*G*^†^ is the height of the free energy barrier and *β* = (*k*_B_*T*)^−1^. Using Δ*G*^†^ ∼ 3.5kT from the PBMetaD simulations and the fact that the height of the barrier is approximately the same on both sides of the transition, we obtain 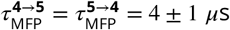.

The analyses above enable us to draw some additional conclusions about the dynamics of the inhibitory module in the inactive state of the complex. Our simulations show that the Ac domain is in a fast equilibrium between a tightly bound state (state **5**) and one were the Ac-helix is more relaxed (state **4**) but still inhibits the function of the DH domain by burying residue Y174. This is coherent with the observation that the the Ac-domain binds to the DH one mostly through hydrophobic interactions and establishes a hydrogen bond only between Y174 and R332. The existence of this fast equilibrium is compatible with experimental results, which suggest the dynamics of the inhibitory module to occur at multiple timescales (***Li et al., 2008***). Ultimately, the fact that the state around that CV1 ∼ 0.95 approximates well a transition state also indicates that CV1 is a good guess for the reaction coordinate that drives the transition from the bound state to the molten globule-like state.

### The binding equilibrium

The free energy landscape (Fig. 2A) provides an overall picture of the binding dynamics of the inhibitory module to the DH domain, which is further corroborated by the unbiased MD simulations (Fig. S8). Previously, the binding of the Ac helix to the DH domain has been studied using two types of NMR experiments (***Li et al., 2008***), and it would be relevant to compare our simulations with these.

First, relaxation dispersion measurements of a number of methyl groups revealed the presence of microsecond–millisecond dynamics at a number of sites in the Vav1 AD complex (***Li et al., 2008***). Residues experiencing such dynamics were located both in the Ac helix and in the Ac helix binding site, but also throughout the rest of the protein. Dispersion measurements on the phosphorylated protein (pY174) showed quenched dynamics in those residues in the Ac helix that displayed exchange dynamics in the unphosphorylated protein. However, extensive dynamics dynamics remained in many other residues. Thus, the dispersion experiments suggest multiple dynamical processes (with at least four states) on the microsecond–millisecond time scales, with only some of these related directly to the binding of the Ac helix to the DH domain. This complicated exchange dynamics prevented quantitative fitting and modelling of the relaxation dispersion NMR data (***Li et al., 2008***).

In a second set of experiments, the authors studied a series of mutant variants of Vav1 that shift the equilibrium between the bound and unbound Ac helix. The NMR measurements show that the binding-unbinding reaction is fast on the NMR timescale and were used to estimate the populations of the unbound state, which for the wild-type protein was estimated to be 9% corresponding to a Δ*G*_exp_ = – 2.3 k_B_T. The suggestion that this equilibrium corresponds to opening of the protein and exposure of Y174 to solvent was supported by a high correlation between the estimated population of the unbound state for the mutants and the rate of phosphorylation by the Lck kinase.

Together, the NMR experiments thus suggests a complex dynamical landscape between multiple states, making it difficult to compare our simulations directly to experiments. In particular, the interpretation of both sets of NMR experiments are dependent on both the rates of conformational exchange as well as the differences in chemical shifts in the different substates. Since both experiments and simulations revealed that the Vav1 dynamics is not a simple two-state system it remains somewhat unclear which structures should be considered part of the bound state and which would be unbound.

We thus proceeded by estimating the population of bound and unbound structures by integrating the populations from the free energy surface along CV1 and CV2 using different definitions of the bound state (from CV1*>* 0.5 to CV1*>* 0.95) and defining the unbound state as those that were not bound (i.e. as a two-state system; ***Lindorff-Larsen and Teilum (2021)***). The resulting free energy differences range from Δ*G*_sim_ = 0.6±0.3 to 3±1 k_B_T (where the uncertainties have been determined via block error analysis) depending on the definition of the bound state (Fig. S10).

These results are in apparent disagreement with the experimentally determined free energy difference between the two states (Δ*G*_exp_ = –2.3 k_B_T) (***Li et al., 2008***). Possible reasons behind this inconsistency include (i) the possible lack of convergence in the estimation of Δ*G*_sim_, which may in part be due to difficulties in overcoming the entropic barrier separating the bound from the unbound states, and (ii) possible imperfections in the force field that we used. These points are discussed below.

Concerning the issue of convergence, we performed a block error analysis (***Flyvbjerg and Petersen, 1989***) to estimate the error of Δ*G*_sim_ (Fig. S11). The monotonously increasing behavior of the the calculated errors for almost every definition of the unbound state suggests that the values are not fully converged. While the difference between experimental and calculated values of Δ*G* are greater than the error, the block analysis suggests that the error estimates themselves might be uncertain. This implies that the discrepancy could, at least partially, be explained by the difficulty in converging this quantity.

One of the reasons that the system is difficult to sample is that binding events may occur through ‘entropic funnels’ that are usually not simply discernible by projecting the dynamics on a small number of collective variables. In the context of the Ac helic binding to the DH domain, there appears to be an entropic barrier separating the unbound state **2** from the partially bound state **3** (Fig. 2). Although such a barrier is not clearly visible in the free energy landscape projected onto CV1 and CV2, a strong indication of the difficulty in observing this transition comes from the fact that the multiple shorter MD simulations are able to interconnect all the metastable states (Fig. 2) except for **2** and **3** (Fig. S8C,D). This makes it particularly hard to observe multiple, independent binding-unbinding events, and even enhanced sampling methods are difficult to apply because of the inherent degeneracy of the unbound state. While methods such as by Funnel Metadynamics (***Limongelli et al., 2013***) can be used to enhance sampling of binding events, they have mostly been employed to relatively rigid ligands.

Finally, lack of agreement between experiment and simulation can be due to inaccuracies in the force field. In our simulations we used the Amber a99SB-*disp* force field (***Robustelli et al., 2018***) as one of the only force fields that has been extensively tested and found to describe both folded and disordered proteins relatively well. This ability is crucial to study systems such as Vav1 that contain both disordered and ordered regions (***Robustelli et al., 2020***). A key aspect of constructing a force field with these properties was to obtain a better balance between protein-protein and protein-water interactions (***Piana et al., 2015***; ***Robustelli et al., 2018***). Nevertheless, it is possible that remaining imbalances in both solvent and other interactions can explain the difference in affinity for the Ac helix. Indeed, a recent analysis has shown that a99SB-*disp* underestimates the stability of several complexes between folded proteins (***Piana et al., 2020***).

As a final remark, we note that a more detailed comparison to the experimental measurements on the Vav1 binding-unbinding equilibrium would require calculating the methyl NMR chemical shifts from the simulations and comparing with those in experiments (***Li et al., 2008***). Such comparisons are, however, complicated by the fact that (i) they would require simulating the protein variants that were used to probe the different Ac-binding equilibria and (ii) the equilibrium populated by the protein encompasses multiple states that are difficult to decouple in experimental data. It would also be possible to calculate the exchange contribution to the transverse relaxation rate either by performing longer simulations (***Xue et al., 2012***; ***Lindorff-Larsen et al., 2016***) or constructing an MSM (***Olsson and Noé, 2017***) and thereby compare directly to these NMR measurements (***Li et al., 2008***).

## Conclusions

Comparisons of phosphoproteomics data with structural information has revealed the intriguing observation that a number of phosphorylations occur at positions that are buried inside the protein structure in the unphosphorylated state (***Jiménez et al., 2007***). One hypothesis for how these sites may become accessible to kinases is that they exist in a dynamic equilibrium between buried and exposed states, and such dynamics have been shown for example for both Vav1 (***Li et al., 2008***) and p27 in complex with its targets (***Tsytlonok et al., 2019***; ***Henriques and Lindorff-Larsen, 2020***). From a functional perspective, these proteins are particularly interesting because both states are essential for function: the major state is an inhibited stated, and the minor state is necessary for the ability of the kinase to relieve inhibition. One would thus expect that the balance between the two states is finely tuned during evolution. From a biophysical perspective, such systems are good model systems to study folding upon binding of intrinsically disordered proteins. Because the disordered region is tethered to the folded region, and because the populations of the bound and the unbound states are known and roughly equal, the system might also be very useful to benchmark force fields.

We have investigated the binding dynamics of the Ac domain to the DH domain and used both standard MD and metadynamics simulations to probe the transient exposure of Y174 in Vav1. Despite the disagreement with NMR experiments in terms of the ratio between the active and the inhibited states populations, we have found a complex binding pathway that proceeds by folding the Ac-helix while unbound and accommodating it in the hydrophobic pocket formed by its binding site. This leads to the conclusion that the binding mechanism involves association between a partially helical structure and the DH domain, which then proceeds via a molten-globule like intermediate and a major transition before reaching the final bound state.

## Methods

### Protein structures

Two structures are available for the AD complex: one solved by NMR (PDB: 1F5X, ***Aghazadeh et al. (2000)***) and one by crystallography (PDB: 3KY9, ***Yu et al. (2010)***). While the NMR structure corresponds to the 168-365 construct, similar to the one used to study dynamics in Vav1 (***Li et al., 2008***), the crystal structure is composed by 5 domains (in a dimeric form) and lacks coordinates for the 181-188 loop. In order to prepare the crystal structure for MD simulations, we isolated the 169-365 fragment of the first monomer and modelled the missing loop using Modeller (***Eswar et al., 2006***). In particular, we generated 50 different models and selected the one with the best Modeller score. In the case of the NMR structure, instead, we removed residue A168 to match the sequence used in the NMR experiments (***Li et al., 2008***).

### Molecular dynamics

We used GROMACS v.2018.6 (***Abraham et al., 2015***) for our MD simulations. The protein structures were solvated in a dodecahedral box of water molecules with periodic boundary conditions. We set the dimension of the box in a such a way that the protein was 1 nm away from any side of the solvent box (*d* = 8.3 nm for the native, autoinhibited state) resulting in approximately 12,000 water molecules. Chloride and sodium ions were included to neutralize the protein and to set the salt concentration to 50 mM. In most of the simulations we employed the a99SB-*disp* force field (***Robustelli et al., 2018***) with its corresponding TIP4PD-like water model, while we conducted some other control simulations using CHARMM36m force field (***Huang et al., 2017***) with the CHARMM-modified TIP3P water model.

The Van der Waals interactions were set to zero at 1.0 and 1.2 nm cutoff, for AMBER and CHARMM respectively, and the long-range electrostatics was taken into account by means of the particle mesh Ewald algorithm with a 0.16 nm mesh spacing and cubic interpolation. The bonds involving hydrogen atoms were constrained using the LINCS algorithm. Equations of motion were integrated using the leapfrog algorithm with a 2 fs time step. We employed the Bussi thermostat (***Bussi et al., 2007***) to control the temperature, with τ_T_ = 0.1 ps^−1^ coupling time constant. Before production, we used steepest descent to minimize the energy, and then equilibrated the box first for 5 ns in the NVT ensemble at 300K and then in the NPT ensemble, employing in both cases position restraints of 1000 kJ mol^−1^ nm^−2^ on all interatomic bonds. We fixed the pressure to 1 atm in the NPT ensemble through the Parrinello-Rahman barostat (***Parrinello and Rahman, 1981***), setting τ_P_ = 2 ps^−1^ as coupling time constant and an isothermal compressibility of 4.5 × 10^−5^ bar^−1^.

### Well-tempered and parallel bias metadynamics

We used the PLUMED v.2.5 plugin (***Tribello et al., 2014***) together with Gromacs to perform both well-tempered and multiple-walker parallel-bias metadynamics (respectively, WTMetaD and PBMetaD) simulations (***Barducci et al., 2008***; ***Pfaendtner and Bonomi, 2015***). In these simulations we used 2–5 collective variables (CVs), described below and in the supplementary material. Gaussians were deposited every 1 ps, with an initial height of 1 kJ/mol, and a bias factor *γ*^−1^ = 10 for WTMetaD and *γ*^−1^ = 44 for PBMetaD. In the case of PBMetaD, we employed two walkers that were simultaneously depositing Gaussians on the five one-dimensional CVs. Input files for PLUMED can be found in PLUMED-NEST (***Bonomi et al., 2019***) at https://www.plumed-nest.org/eggs/21/017/. The initial configurations for metadynamics simulation were prepared as described above apart from the box size. To avoid self-interaction across the periodic boundaries in the unbound, active state (which is substantially more extended than the inhibited state), we adopted the following strategy. We first performed a (non well-tempered) metadynamics simulation with Gaussian hills heights of 3 kJ/mol in order to induce a rapid transition to an extended state of the Ac-helix. We let the simulation run for 50 ns and selected the frame corresponding to the most extended configuration, which was then re-solvated. This resulted in a substantially larger box (*d* = 11.4 nm) with respect to the one used for standard MD simulations described above (34,000 water molecules and a total of 130,000 particles). We then solvated the bound (crystal) structure in a box of this size and used this as starting point for the WTMetaD. This structure was also used as starting point for one of the replicas in the PBMetaD simulations, and the other replica was started from an unbound structure extracted from the WTMetaD simulation.

### Analysis

All analyses were conducted using GROMACS, PLUMED, VMD (***Humphrey et al., 1996***), MDTraj (***McGibbon et al., 2015***) and in-house developed Python tools using Numpy (***Harris et al., 2020***), Scipy (***Virtanen et al., 2020***) and PyEMMA (***Scherer et al., 2015***); see https://github.com/KULL-Centre/papers/tree/main/2021/Vav1-Orioli-et-al for the simulations, data, supplementary videos and the scripts employed to generate the paper figures and https://github.com/fpesceKU/BLOCKING for the code employed to perform block error analysis. We obtained free energy surfaces for WT-MetaD by employing the sum_hills tool in PLUMED, while we reweighted PBMetaD simulations following the prescription in ***Pfaendtner and Bonomi (2015)*** after discarding the first 1 *µ*s as equilibration (20% of the simulation length) from both the walkers. All plots were made using Matplotlib (***Hunter, 2007***), while all the renderings and supplementary videos were obtained using VMD and the Tachyon software (***Stone et al., 2013***).

## Acknowledgments

We acknowledge work by Fabio Doro who contributed to the early stages and simulations in this project, and to Micha B. A. Kunze and Elena Papaleo for fruitful discussions. We are grateful to Francesco Pesce for inspiring discussions and for providing his scripts to perform block error analysis. We acknowledge support from the NordForsk Nordic Neutron Science Programme (grant number 81912), the Lundbeck Foundation BRAINSTRUC structural biology initiative (R155-2015-2666) and Independent Research Fund Denmark (grant number 6108-00471). We also acknowledge access to computational resources from the ROBUST Resource for Biomolecular Simulations (supported by the Novo Nordisk Foundation grant no. NF18OC0032608), the Danish National Supercomputer for Life Sciences (Computerome) and the Biocomputing Core Facility at the Department of Biology, University of Copenhagen.

## References

Abe K, Whitehead IP, O’Bryan JP, Der CJ. Involvement of NH2-terminal sequences in the negative regulation of Vav signaling and transforming activity. Journal of Biological Chemistry. 1999; 274(43):30410–30418.

Abraham MJ, Murtola T, Schulz R, Páll S, Smith JC, Hess B, Lindahl E. GROMACS: High performance molecular simulations through multi-level parallelism from laptops to supercomputers. SoftwareX. 2015; 1:19–25.

Aghazadeh B, Lowry WE, Huang XY, Rosen MK. Structural basis for relief of autoinhibition of the Dbl homology domain of proto-oncogene Vav by tyrosine phosphorylation. Cell. 2000; 102(5):625–633.

Barducci A, Bussi G, Parrinello M. Well-tempered metadynamics: a smoothly converging and tunable freeenergy method. Physical review letters. 2008; 100(2):020603.

Best RB, Hummer G, Eaton WA. Native contacts determine protein folding mechanisms in atomistic simulations. Proceedings of the National Academy of Sciences. 2013; 110(44):17874–17879.

Bonomi M, Bussi G, Camilloni C, Tribello GA, Banás P, Barducci A, Bernetti M, Bolhuis PG, Bottaro S, Branduardi D, et al. Promoting transparency and reproducibility in enhanced molecular simulations. Nature methods. 2019; 16(8):670–673.

Bussi G, Donadio D, Parrinello M. Canonical sampling through velocity rescaling. The Journal of chemical physics. 2007; 126(1):014101.

Bussi G, Laio A. Using metadynamics to explore complex free-energy landscapes. Nature Reviews Physics. 2020; p. 1–13.

Bussi G, Tribello GA. Analyzing and biasing simulations with PLUMED. In: Biomolecular Simulations Springer; 2019. p. 529–578.

Bustelo XR. Vav proteins, adaptors and cell signaling. Oncogene. 2001; 20(44):6372–6381.

Chodera JD, Noé F. Markov state models of biomolecular conformational dynamics. Current opinion in structural biology. 2014; 25:135–144.

Chung HS, Louis JM, Eaton WA. Experimental determination of upper bound for transition path times in protein folding from single-molecule photon-by-photon trajectories. Proceedings of the National Academy of Sciences. 2009; 106(29):11837–11844.

Dombroski AJ, Walter WA, Gross CA. Amino-terminal amino acids modulate sigma-factor DNA-binding activity. Genes & development. 1993; 7(12a):2446–2455.

Du R, Pande VS, Grosberg AY, Tanaka T, Shakhnovich ES. On the transition coordinate for protein folding. The Journal of chemical physics. 1998; 108(1):334–350.

Dyson HJ, Wright PE. Coupling of folding and binding for unstructured proteins. Current opinion in structural biology. 2002; 12(1):54–60.

Eswar N, Webb B, Marti-Renom MA, Madhusudhan M, Eramian D, Shen My, Pieper U, Sali A. Comparative protein structure modeling using Modeller. Current protocols in bioinformatics. 2006; 15(1):5–6.

Flyvbjerg H, Petersen HG. Error estimates on averages of correlated data. The Journal of Chemical Physics. 1989; 91(1):461–466.

Fu CD, Pfaendtner J. Lifting the curse of dimensionality on enhanced sampling of reaction networks with parallel bias metadynamics. Journal of chemical theory and computation. 2018; 14(5):2516–2525.

Habchi J, Tompa P, Longhi S, Uversky VN. Introducing protein intrinsic disorder. Chemical reviews. 2014; 114(13):6561–6588.

Harris CR, Millman KJ, van der Walt SJ, Gommers R, Virtanen P, Cournapeau D, Wieser E, Taylor J, Berg S, Smith NJ, Kern R, Picus M, Hoyer S, van Kerkwijk MH, Brett M, Haldane A, del R’io JF, Wiebe M, Peterson P, G’erard-Marchant P, et al. Array programming with NumPy. Nature. 2020 Sep; 585(7825):357–362. https://doi.org/10.1038/s41586-020-2649-2, doi: 10.1038/s41586-020-2649-2.

Heller GT, Aprile FA, Bonomi M, Camilloni C, De Simone A, Vendruscolo M. Sequence specificity in the entropy-driven binding of a small molecule and a disordered peptide. Journal of molecular biology. 2017; 429(18):2772–2779.

Henriques J, Lindorff-Larsen K. Protein dynamics enables phosphorylation of buried residues in Cdk2/Cyclin A-bound p27. bioRxiv. 2020;.

Huang J, Rauscher S, Nawrocki G, Ran T, Feig M, de Groot BL, Grubmüller H, MacKerell AD. CHARMM36m: an improved force field for folded and intrinsically disordered proteins. Nature methods. 2017; 14(1):71–73.

Hubbard SR. Juxtamembrane autoinhibition in receptor tyrosine kinases. Nature Reviews Molecular Cell Biology. 2004; 5(6):464–471.

Hummer G. From transition paths to transition states and rate coefficients. The Journal of chemical physics. 2004; 120(2):516–523.

Humphrey W, Dalke A, Schulten K, et al. VMD: visual molecular dynamics. Journal of molecular graphics. 1996; 14(1):33–38.

Hunter JD. Matplotlib: A 2D graphics environment. Computing in Science & Engineering. 2007; 9(3):90–95. doi: 10.1109/MCSE.2007.55.

Jiménez JL, Hegemann B, Hutchins JR, Peters JM, Durbin R. A systematic comparative and structural analysis of protein phosphorylation sites based on the mtcPTM database. Genome biology. 2007; 8(5):1–20.

Kim AS, Kakalis LT, Abdul-Manan N, Liu GA, Rosen MK. Autoinhibition and activation mechanisms of the Wiskott–Aldrich syndrome protein. Nature. 2000; 404(6774):151–158.

Kim JY, Meng F, Yoo J, Chung HS. Diffusion-limited association of disordered protein by non-native electrostatic interactions. Nature communications. 2018; 9(1):4707.

Ko LJ, Prives C. p53: puzzle and paradigm. Genes & development. 1996; 10(9):1054–1072.

Lee GM, Donaldson LW, Pufall MA, Kang HS, Pot I, Graves BJ, McIntosh LP. The structural and dynamic basis of Ets-1 DNA binding autoinhibition. Journal of Biological Chemistry. 2005; 280(8):7088–7099.

Li P, Martins IR, Amarasinghe GK, Rosen MK. Internal dynamics control activation and activity of the autoinhibited Vav DH domain. Nature structural & molecular biology. 2008; 15(6):613–618.

Limongelli V, Bonomi M, Parrinello M. Funnel metadynamics as accurate binding free-energy method. Proceedings of the National Academy of Sciences. 2013; 110(16):6358–6363.

Lindorff-Larsen K, Teilum K. Linking thermodynamics and measurements of protein stability. Protein Eng Des Sel. 2021 Feb; 34.

Lindorff-Larsen K, Maragakis P, Piana S, Shaw DE. Picosecond to millisecond structural dynamics in human ubiquitin. The Journal of Physical Chemistry B. 2016; 120(33):8313–8320.

Malagrinò F, Visconti L, Pagano L, Toto A, Troilo F, Gianni S. Understanding the Binding Induced Folding of Intrinsically Disordered Proteins by Protein Engineering: Caveats and Pitfalls. International Journal of Molecular Sciences. 2020; 21(10):3484.

McGibbon RT, Beauchamp KA, Harrigan MP, Klein C, Swails JM, Hernández CX, Schwantes CR, Wang LP, Lane TJ, Pande VS. MDTraj: A Modern Open Library for the Analysis of Molecular Dynamics Trajectories. Biophysical Journal. 2015; 109(8):1528–1532. doi: 10.1016/j.bpj.2015.08.015.

Noé F, Wu H, Prinz JH, Plattner N. Projected and hidden Markov models for calculating kinetics and metastable states of complex molecules. The Journal of chemical physics. 2013; 139(18):11B609_1.

Olsson S, Noé F. Mechanistic models of chemical exchange induced relaxation in protein NMR. Journal of the American Chemical Society. 2017; 139(1):200–210.

Orioli S, Faccioli P. Dimensional reduction of Markov state models from renormalization group theory. The Journal of chemical physics. 2016; 145(12):124120.

Papaleo E, Sutto L, Gervasio FL, Lindorff-Larsen K. Conformational changes and free energies in a proline isomerase. Journal of Chemical Theory and Computation. 2014; 10(9):4169–4174.

Parrinello M, Rahman A. Polymorphic transitions in single crystals: A new molecular dynamics method. Journal of Applied physics. 1981; 52(12):7182–7190.

Pearson MA, Reczek D, Bretscher A, Karplus PA. Structure of the ERM protein moesin reveals the FERM domain fold masked by an extended actin binding tail domain. Cell. 2000; 101(3):259–270.

Pérez-Hernández G, Paul F, Giorgino T, De Fabritiis G, Noé F. Identification of slow molecular order parameters for Markov model construction. The Journal of chemical physics. 2013; 139(1):07B604_1.

Pfaendtner J, Bonomi M. Efficient sampling of high-dimensional free-energy landscapes with parallel bias metadynamics. Journal of chemical theory and computation. 2015; 11(11):5062–5067.

Piana S, Donchev AG, Robustelli P, Shaw DE. Water dispersion interactions strongly influence simulated structural properties of disordered protein states. The journal of physical chemistry B. 2015; 119(16):5113–5123.

Piana S, Lindorff-Larsen K, Shaw DE. Atomistic description of the folding of a dimeric protein. The Journal of Physical Chemistry B. 2013; 117(42):12935–12942.

Piana S, Robustelli P, Tan D, Chen S, Shaw DE. Development of a force field for the simulation of single-chain proteins and protein–protein complexes. Journal of chemical theory and computation. 2020; 16(4):2494–2507.

Pietrucci F, Laio A. A collective variable for the efficient exploration of protein beta-sheet structures: application to SH3 and GB1. Journal of Chemical Theory and Computation. 2009; 5(9):2197–2201.

Prakash A, Sprenger K, Pfaendtner J. Essential slow degrees of freedom in protein-surface simulations: A metadynamics investigation. Biochemical and biophysical research communications. 2018; 498(2):274–281.

Pufall MA, Graves BJ. Autoinhibitory domains: modular effectors of cellular regulation. Annual review of cell and developmental biology. 2002; 18(1):421–462.

Pufall MA, Lee GM, Nelson ML, Kang HS, Velyvis A, Kay LE, McIntosh LP, Graves BJ. Variable control of Ets-1 DNA binding by multiple phosphates in an unstructured region. science. 2005; 309(5731):142–145.

Röblitz S, Weber M. Fuzzy spectral clustering by PCCA+: application to Markov state models and data classification. Advances in Data Analysis and Classification. 2013; 7(2):147–179.

Robustelli P, Piana S, Shaw DE. Developing a molecular dynamics force field for both folded and disordered protein states. Proceedings of the National Academy of Sciences. 2018; 115(21):E4758–E4766.

Robustelli P, Piana S, Shaw DE. The mechanism of coupled folding-upon-binding of an intrinsically disordered protein. Journal of the American Chemical Society. 2020;.

Scherer MK, Trendelkamp-Schroer B, Paul F, Pérez-Hernández G, Hoffmann M, Plattner N, Wehmeyer C, Prinz JH, Noé F. PyEMMA 2: A Software Package for Estimation, Validation, and Analysis of Markov Models. Journal of Chemical Theory and Computation. 2015 Oct; 11:5525–5542. http://dx.doi.org/10.1021/acs.jctc.5b00743, doi: 10.1021/acs.jctc.5b00743.

Schuler B, Borgia A, Borgia MB, Heidarsson PO, Holmstrom ED, Nettels D, Sottini A. Binding without folding–the biomolecular function of disordered polyelectrolyte complexes. Current opinion in structural biology. 2020; 60:66–76.

Schwantes CR, Pande VS. Improvements in Markov state model construction reveal many non-native interactions in the folding of NTL9. Journal of chemical theory and computation. 2013; 9(4):2000–2009.

Sgourakis NG, Patel MM, Garcia AE, Makhatadze GI, McCallum SA. Conformational dynamics and structural plasticity play critical roles in the ubiquitin recognition of a UIM domain. Journal of molecular biology. 2010; 396(4):1128–1144.

Sidky H, Chen W, Ferguson AL. High-resolution Markov state models for the dynamics of Trp-cage miniprotein constructed over slow folding modes identified by state-free reversible VAMPnets. The Journal of Physical Chemistry B. 2019; 123(38):7999–8009.

Stone JE, Vandivort KL, Schulten K. GPU-accelerated molecular visualization on petascale supercomputing platforms. In: Proceedings of the 8th International Workshop on Ultrascale Visualization; 2013. p. 1–8.

Stöven S, Ando I, Kadalayil L, Engström Y, Hultmark D. Activation of the Drosophila NF-kappaB factor Relish by rapid endoproteolytic cleavage. EMBO Rep. 2000; 1(4):347–352.

Sturzenegger F, Zosel F, Holmstrom ED, Buholzer KJ, Makarov DE, Nettels D, Schuler B. Transition path times of coupled folding and binding reveal the formation of an encounter complex. Nature communications. 2018; 9(1):4708.

Tribello GA, Bonomi M, Branduardi D, Camilloni C, Bussi G. PLUMED 2: New feathers for an old bird. Computer Physics Communications. 2014; 185(2):604–613.

Trudeau T, Nassar R, Cumberworth A, Wong ET, Woollard G, Gsponer J. Structure and intrinsic disorder in protein autoinhibition. Structure. 2013; 21(3):332–341.

Tsytlonok M, Sanabria H, Wang Y, Felekyan S, Hemmen K, Phillips AH, Yun MK, Waddell MB, Park CG, Vaithiyalingam S, et al. Dynamic anticipation by Cdk2/Cyclin A-bound p27 mediates signal integration in cell cycle regulation. Nature communications. 2019; 10(1):1–13.

Tybulewicz VL. Vav-family proteins in T-cell signalling. Current opinion in immunology. 2005; 17(3):267–274.

Van Der Lee R, Buljan M, Lang B, Weatheritt RJ, Daughdrill GW, Dunker AK, Fuxreiter M, Gough J, Gsponer J, Jones DT, et al. Classification of intrinsically disordered regions and proteins. Chemical reviews. 2014; 114(13):6589–6631.

Virtanen P, Gommers R, Oliphant TE, Haberland M, Reddy T, Cournapeau D, Burovski E, Peterson P, Weckesser W, Bright J, van der Walt SJ, Brett M, Wilson J, Millman KJ, Mayorov N, Nelson ARJ, Jones E, Kern R, Larson E, Carey CJ, et al. SciPy 1.0: Fundamental Algorithms for Scientific Computing in Python. Nature Methods. 2020; 17:261–272. doi: 10.1038/s41592-019-0686-2.

Wang Y, Papaleo E, Lindorff-Larsen K. Mapping transiently formed and sparsely populated conformations on a complex energy landscape. Elife. 2016; 5:e17505.

Xue Y, Ward JM, Yuwen T, Podkorytov IS, Skrynnikov NR. Microsecond time-scale conformational exchange in proteins: using long molecular dynamics trajectory to simulate NMR relaxation dispersion data. Journal of the American Chemical Society. 2012; 134(5):2555–2562.

Yu B, Martins IR, Li P, Amarasinghe GK, Umetani J, Fernandez-Zapico ME, Billadeau DD, Machius M, Tomchick DR, Rosen MK. Structural and energetic mechanisms of cooperative autoinhibition and activation of Vav1. Cell. 2010; 140(2):246–256.

