## Supporting Text, Figures and Table for "Transient exposure of a buried phosphorylation site in an autoinhibited protein"

### Supporting Information

#### Collective Variables

We used one or more of five different CVs in this manuscript.

The first, CV1, quantifies the extent to which the Ac-helix is bound in its native position to the DH-domain. Specifically, we define CV1 via a continuous function quantifying the fraction of a set of native contacts that are formed (***Best et al., 2013***):

$$\mathrm{CV1}(X) = \sum_{i \in S_1} \sum_{j \in S_2} \frac{1}{1 + \exp\left[\beta \left(r_{ij}(X) - \lambda r_{ij}(X_0)\right)\right]} \tag{2}$$

Here, $X$ represents the instantaneous configuration of the protein, $S_1$ and $S_2$ are two sets of atoms, $r_{ij}(X)$ is the distance between atoms $i$ and $j$ in configuration X, $r^0_{ij}$ is the distance between the same two atoms in a reference conformation $X_0$, $\beta = 5$Å$^{-1}$ and $\lambda = 2$. Set $S_1$ comprises of all the heavy atoms in the Ac-helix (residue 169 to 178), while set $S_2$ is defined by all the heavy atoms $j$ in a reference structure found closer than 6Å from any atom $i \in S_1$ such that the amino acids to which $i$ and $j$ belong are at least 3 residues apart in the protein's sequence. In the the simulations started from the NMR structure, we used this structure as the reference for defining the CVs, and similarly used crystal structure as reference conformation for the simulations started from that structure. With this choice, some intra-helix contacts are included in the definition of CV1 and this is useful to distinguish between unbound states of the helix with folded helix (CV1$\sim 0.25$) and unbound states with unfolded helix (CV1$\sim 0$).

The second collective variable CV2, instead, measures the amount of helical structure in the 169-178 portion of the Ac domain. This is achieved by measuring the cosine distance between the instantaneous value of the dihedral angles in the Ac-helix with those of a reference structure $X_0$:

$$\mathrm{CV2}(X) = \frac{1}{2} \sum_{i=1}^{N} \left[1 + \cos(\phi(X) - \phi(X_0))\right] \tag{3}$$

The reference structure for CV2 is the same as for CV1.

The third collective variable, CV3, is defined identically to CV1 but with different choices for the atomic sets $S_1$ and $S_2$. In particular, $S_1$ contains the heavy atoms in the side chain for Y174 and the corresponding $C_\alpha$, while $S_2$ is defined by all the heavy atoms in the sidechains of T205, Y209, V328, Q331 and R332. Also in the case of $S_2$, the alpha-carbons of each residue have been included as well. In this way, CV3 can be interpreted as a measure of the binding of Y174 to its binding site, irregardless of the conformation of the Ac-helix. Similarly, CV4 is defined as the fraction of contacts between the alpha-carbon and sidechain heavy atoms in I173, Y174 and L177 and those in Y209, T212, P320, L325 and V328. Despite its similarity with CV3, CV4 introduces contacts relevant for the appropriate positioning of the Ac-helix, especially at the beginning of the binding process.

Finally, the last collective variable CV5 is based on ***Pietrucci and Laio*** (***2009***) who introduced a different way to estimate the helical content. In particular, we used

$$\mathrm{CV5}(X) = \sum_{i=1}^{4} \frac{1 - \left(\frac{r_i}{r_0}\right)^6}{1 - \left(\frac{r_i}{r_0}\right)^1 2} \tag{4}$$

where $r_i$ measure the distances from an alpha-helical configuration (for residues 169–178), $r_0 = 0.1$ nm and the sum runs over the four 6-residue long stretches of the helix.

#### Differences between simulations started from the NMR and crystal structures

The two protein structures, respectively resolved through NMR and crystallography and here labelled as *NMR* and *Crystal*, show clear differences in particular in two regions: (i) In the NMR structure the docking pose of the Ac-helix is almost perfectly horizontal with respect to the DH domain, while in the case of the crystal the acidic domain is slightly tilted, with the N-terminus pointing downwards; (ii) Residues 191–196 show a kink that splits the corresponding helix in two parts, while such a kink is absent in the crystal structure. These structural differences also affect the side chain packing of Y174, which forms a seemingly stable hydrogen bond with R332 in the crystal structure. The same interaction is not present in the NMR structure. We performed 1 $\mu$s-long simulations starting from either the crystal or the NMR structure and using two different force fields (a99sb-*disp* and CHARMM36m). In Fig. S1 we show the behavior of the fraction of native contacts with respect to the first simulation frame (Q), RMSD to initial frame (RMSD), CV1, CV2 and CV3 on the different MD simulations. Examining all of these metrics in the simulations obtained with both the force fields, we find that the NMR structure is less stable than the crystal structure on the $\mu$s timescale. Indeed, the NMR structure looses approximately the 20% of its overall contacts within 500 ns of simulation and the binding of the Ac-helix in the simulation started from the NMR structure appears to be less stable than the one started from the crystal configuration ($\langle \mathrm{CV1}_{\mathrm{CRYSTAL}} \rangle = 0.96 \pm 0.02$ and $\langle \mathrm{CV1}_{\mathrm{NMR}} \rangle = 0.83 \pm 0.02$ for AMBER and $\langle \mathrm{CV1}_{\mathrm{CRYSTAL}} \rangle = 0.95 \pm 0.02$ and $\langle \mathrm{CV1}_{\mathrm{NMR}} \rangle = 0.71 \pm 0.02$ for CHARMM). This suggests that a substantial rearrangement of the interface between the DH and the Ac domains occurs in a short timescale in the case of the simulation started from the NMR structure, as also underlined by the significant change in the packing of Y174 during the simulation. Moreover, we stress the fact that the hydrogen bond between Y173 and R332 is transiently broken and reformed during both the a99SB-*disp* and CHARMM36m simulations started from the crystal structure (Fig. S1), while it is not formed in the runs started from the NMR structure. All these differences are evidently due to a convergence issue, as with infinite sampling the two structures should provide the same equilibrium description. However, the collective variables we employ for biasing need a reference template in order to be properly defined, and for the these reasons we decided to proceed with the crystal structure rather than the NMR structure.

#### Estimation of uncertainties in the calculation of the transition path time

In order to estimate the uncertainty on the calculation of the transition path time and on the committor value we adopted a leave-1-out approach. We split our set of simulations in 5 subsets of 20 simulations each, by removing each time the 5 trajectories started from one of the five selected starting points. Then we computed the value of the binding probability and the transition path time from each of the subsets and averaged the results. Finally, we estimated the uncertainty as the standard error on the sample mean.

#### Definition of states in the PBMetaD free energy landscape

We selected the 5 states along the possible Ac-DH binding pathway by visual inspection of the PBMetaD free energy landscape along CV1 and CV2. In particular, approximating the states as rectangles and using the notation **state** $= (x_0, x_1, y_0, y_1)$, we have:

$$\begin{aligned}
\mathbf{1} &= (0.03, 0.12, 7.5, 10)\\
\mathbf{2} &= (0.175, 0.225, 12.1, 14.2)\\
\mathbf{3} &= (0.56, 0.66, 12.8, 14.3)\\
\mathbf{4} &= (0.75, 0.83, 12.8, 15.55)\\
\mathbf{5} &= (0.95, 1, 13.5, 16)
\end{aligned} \tag{5}$$

The conformations shown in Fig. 2 and 3 have been extracted from the five states among the ones having the highest metadynamics weights.

#### List of the simulations

Here we list all the unbiased MD simulations we conducted to qualitatively assess the shape of the free energy landscape along CV1 and CV2.

| **Starting point** | **# simulations** | **Length** [$\mu$s] |
|---|---|---|
| CV1 = 0.07, CV2 = 9.81 | 3 | 0.5 |
| CV1 = 0.10, CV2 = 8.52 | 2 | 0.5 |
| CV1 = 0.18, CV2 = 13.47 | 3 | 0.5 |
| CV1 = 0.20, CV2 = 12.04 | 4 | 0.5 |
| CV1 = 0.34, CV2 = 15.38 | 5 | 0.5 |
| CV1 = 0.43, CV2 = 14.22 | 5 | 0.3 |
| CV1 = 0.44, CV2 = 13.63 | 1 | 0.5 |
| CV1 = 0.50, CV2 = 14.48 | 6 | 0.3 |
| CV1 = 0.53, CV2 = 13.74 | 2 | 0.5 |
| CV1 = 0.53, CV2 = 13.74 | 2 | 0.5 |
| CV1 = 0.60, CV2 = 13.20 | 1 | 0.5 |
| CV1 = 0.61, CV2 = 14.08 | 2 | 0.5 |
| CV1 = 0.77, CV2 = 13.53 | 3 | 0.5 |
| CV1 = 0.77, CV2 = 15.19 | 1 | 0.5 |
| CV1 = 0.79, CV2 = 13.53 | 3 | 0.5 |
| CV1 = 0.81, CV2 = 14.16 | 1 | 0.5 |
| CV1 = 0.81, CV2 = 15.32 | 1 | 0.5 |
| CV1 = 0.82, CV2 = 12.02 | 1 | 0.5 |
| CV1 = 0.97, CV2 = 14.20 | 4 | 0.3 |
| CV1 = 0.97, CV2 = 15.70 | 1 | 0.5 |
| CV1 = 0.98, CV2 = 15.34 | 1 | 0.5 |

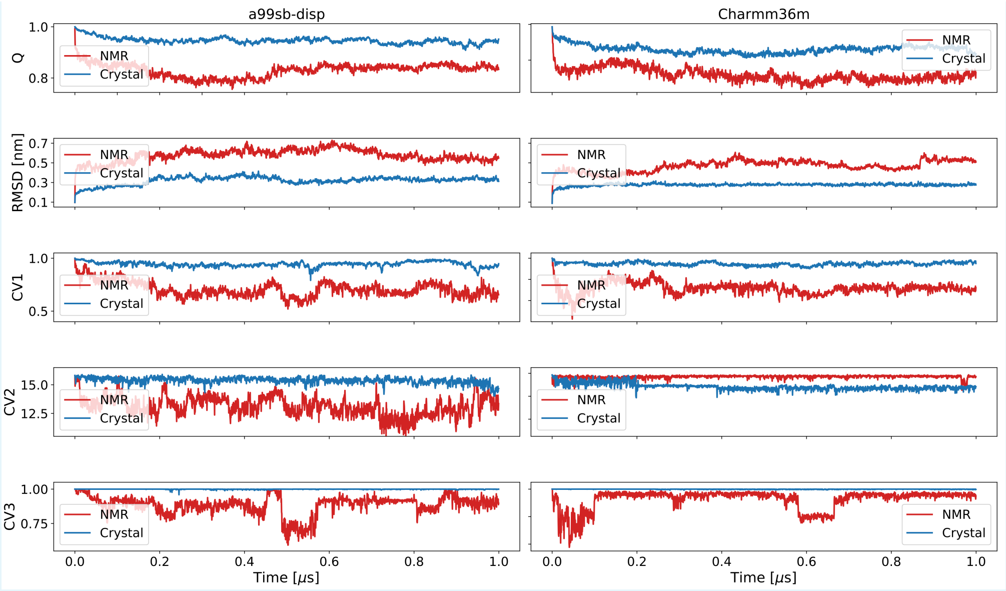


**Supporting Figure 1.** Comparison between the 1 $\mu$s MD simulation of the NMR (red solid line) and crystal (blue solid line) Vav1 AD construct using two different force fields: a99SB-*disp* (left) and CHARMM36m (right).

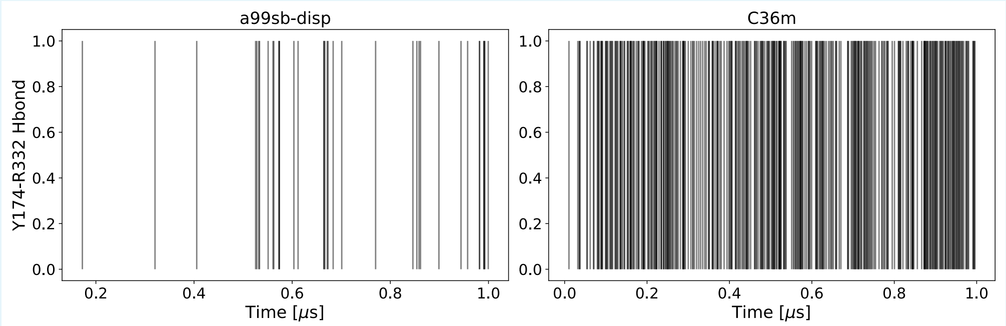


**Supporting Figure 2.** Behaviour of the Y174-R332 hydrogen bond during the MD simulations using, respectively, the a99SB-*disp* and the CHARMM36m force fields started from the crystal configuration of the AD construct. The solid vertical black lines show the time points where the hydrogen bond is formed.

#### WTMetaD simulation

We performed WTMetaD simulation with a bias along CV1 and CV2 only. During the approximately 10 $\mu$s of simulation, we were able to sample multiple binding and unbinding events, both for the whole helix (CV1, black line in Fig. S3 (a)) and for Y174 (Fig. S3B). However, after the potential reached a stationary state, i.e. approximately around 5.5 $\mu$s (see the Gaussian heights in Fig. S3A) the Ac-helix was not able to rebind properly anymore until the very end of the simulation. This is usually a symptom of poor convergence, and indeed the free energy landscape obtained by summing the Gaussian hills (Fig. S3D) confirms this expectation. The free energy surface along CV1 and CV2 shows a sharp minimum around CV1$\sim 0.4$ and CV2$\sim 10$, which appears to be an artefact introduced by the fact that the protein mostly explored that region of CV space during the last 4 $\mu$s of simulation.

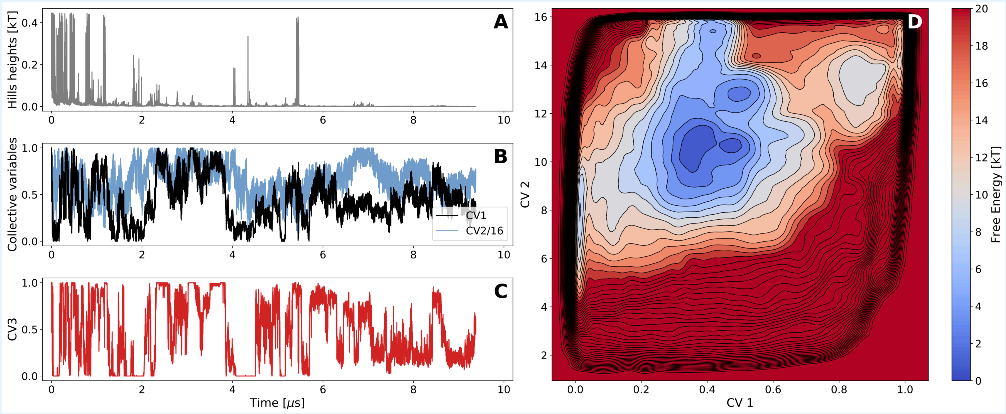


**Supporting Figure 3.** Time evolution of the (A) hills heights, (B) CV1 and CV2 and (C) CV3 in the WTMetaD simulation. (D) Final free energy surface along CV1 and CV2 obtained from the WTMetaD simulation.

The occurrence of such artefacts is often caused by one or more of these issues: (i) the choice of the bias factor $\gamma$ is not optimal and, as a consequence, the bias potential reaches a stationary state before the exploration of the free energy space has been completed; (ii) the choice of the biasing collective variables is not optimal; (iii) the force field is not accurate enough and, when tested on complex conformational transitions and long simulations, it prefers configurations which are not biologically relevant. None of these issues can be easily solved, as both the correct CVs and the correct bias factor can be known precisely only if, respectively, the free energy barriers and the dynamical process under study are known a priori. To mitigate this problem, in the main text we have employed Parallel Bias Metadynamics with 5 collective variables and we have consequently increased the value of the bias factor from 10 to 44.

#### Further details on PBMetaD simulations

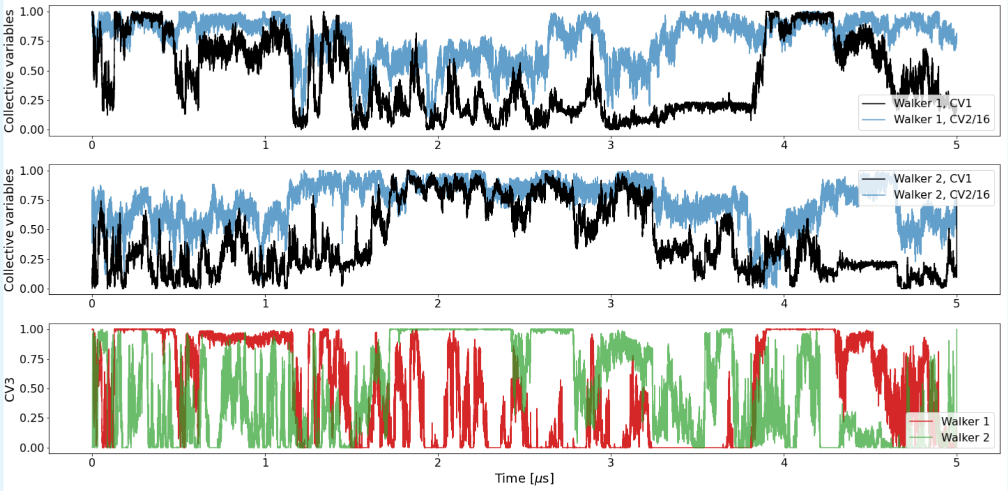


**Supporting Figure 4.** PBMetaD simulations of reversible binding of the Ac helix to the Vav1 DH domain. Dynamics of CV1 and CV2 the (A) first and (B) second walkers. (C) Dynamics of CV3 for both walkers. CV1 and CV2 describe, respectively, the fraction of native contacts between the Ac-helix and the DH domain and the helical content of the Ac helix, while CV3 measures the fraction of native contacts between Y174 and the surrounding residues within a 6Å distance. Values of CV2 have been divided by 16 to bring them onto the same scale as CV1.

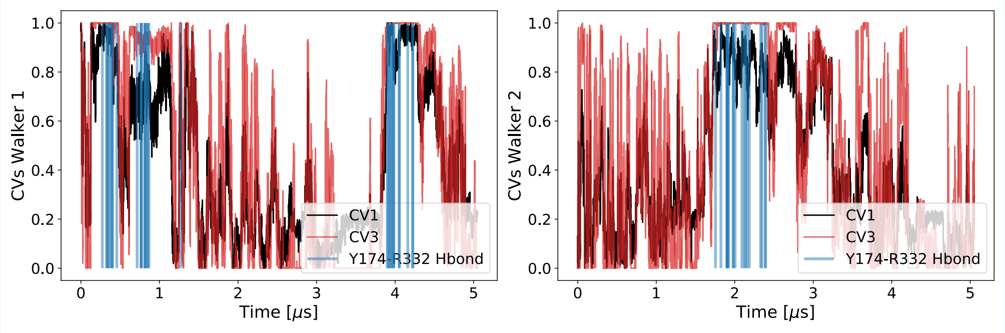


**Supporting Figure 5.** Behaviour of the Y174-R332 hydrogen bond during the PBMetaD simulation. The solid black and red lines show, respectively, the time behaviour of CV1 and CV3 while the blue vertical lines indicate the time points where the hydrogen bond is formed in the simulation.

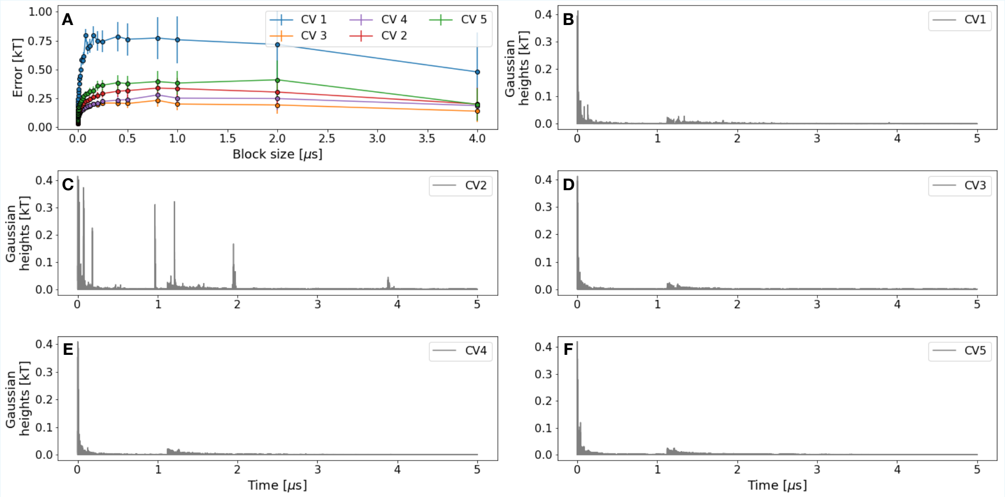


**Supporting Figure 6.** (A) Block error estimates for all CVs used in the PBMetaD simulation; (B)–(F) Time evolution of the Gaussian heights during the PBMetaD simulation for, respectively, CVs from 1 to 5.

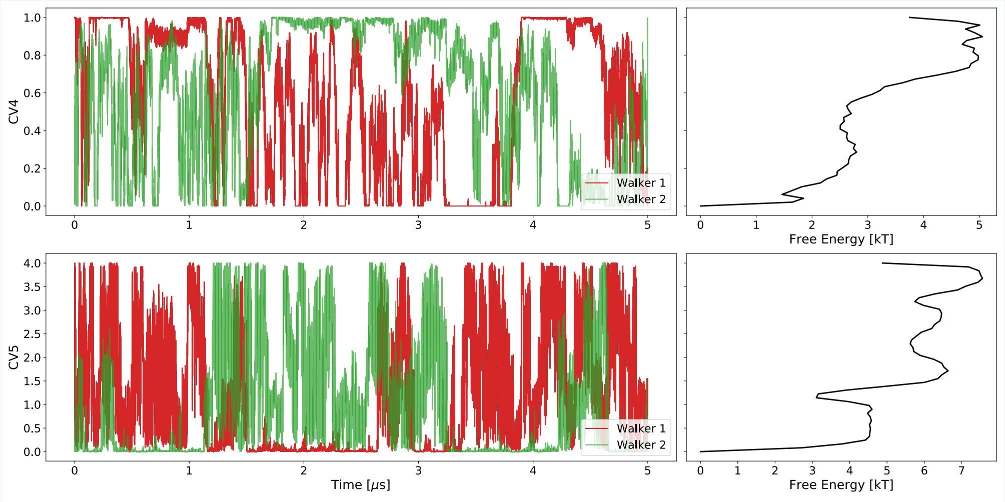


**Supporting Figure 7.** Time evolution of CV4 and CV5 for both walkers in the PBMetaD simulations and the corresponding free energies after reweighting.

#### Unbiased MD simulations

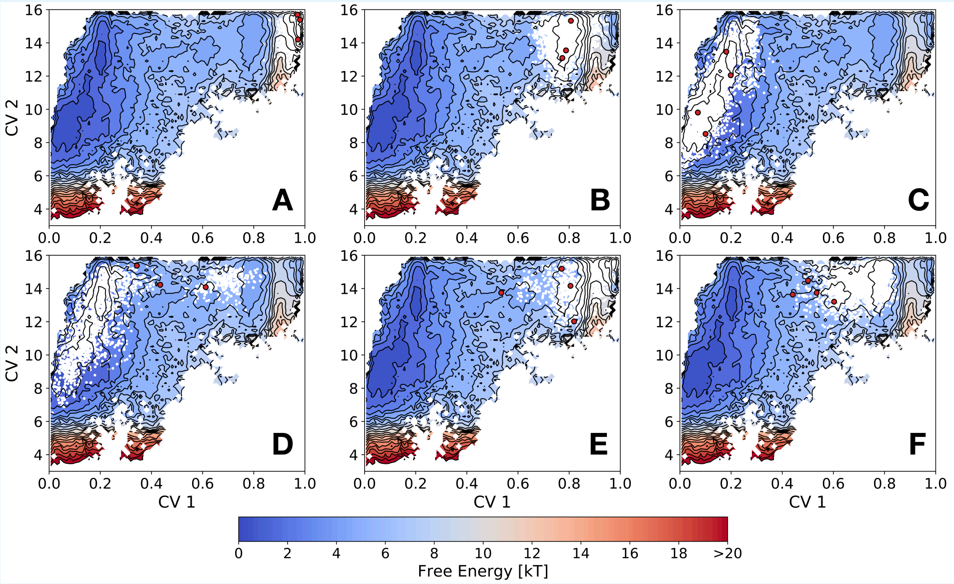


**Supporting Figure 8.** Conformational sampling by unbiased MD simulations initiated from different conformations. We represented the starting point of the simulations as red dots and the conformations sampled in the simulations using smaller white dots. In this way, we aim to highlight the overall regions that are sampled (visualized on the free energy landscape from PBMetaD and defined by CV1 and CV2), rather than visualize individual trajectories. The MD simulations are grouped as follows: (A) simulations sampling the (A) bound state, (B) the molten globule-like state or the (C) unbound state; simulations showing a transition towards the (D) unbound state, (E) bound state or (F) the molten globule state from different locations along the free energy landscape.

#### Constructing a Markov State Model

Before going into the details of the construction of the Markov State Model (MSM), we note that we realized it would be difficult to construct a robust MSM from the sampling that we generated. Assuming that the basins that we observed in the free energy landscapes generated by metadynamics would represent metastable states, we noted that in our unbiased simulations we were not able to connect states **2** and **3** via transitions in both directions. If these states reasonably represent the relevant kinetic states, this evidently introduces a substantial problem in the construction of the MSM, where it is assumed that enough interconnections between all the system's microstates have been observed.

Nevertheless, in an attempt to construct a MSM from our unbiased MD simulations we proceeded as follows. First, we selected as features the inverse distances between the $C_\alpha$ and $C_\beta$ atoms of residues in the Ac-helix with those in the residues in the binding site: all $C_\alpha$ and $C_\beta$ atoms within 6Å from any atom in Ac-helix were considered in the featurization process. Subsequently, we used the VAMP2 score (***Sidky et al., 2019***) as a metric to determine the best lag-time to be employed for time-lagged independent component analysis (TICA) (***Schwantes and Pande, 2013***; ***Pérez-Hernández et al., 2013***), which resulted to be 0.5 ns. When retaining 95% of the cumulative kinetic variance, the TICA was used to reduce the dimension of the initial feature space to ten. Subsequently, we performed clustering in TICA space using the K-Means algorithm, using the VAMP2 score as a heuristic to determine the number of clusters, suggesting 120 as the optimal number. We finally estimated our MSM and computed the first ten implied timescales. The longest implied timescale ($t_1$, Fig. S9A) reports on a relaxation occurring on a timescale of the order of milliseconds, which we can evidently trace back to the bound-to-unbound transition of the Ac-helix, in line with the slow motions observed in the NMR experiments (***Li et al., 2008***). The large separation between the first and remaining timescales suggests that a coarse-grained two state dynamics could be appropriate in the description of the system (***Orioli and Faccioli, 2016***). This is also confirmed by visual inspection of the reweighted free energy (Fig. S9B), which shows a large and degenerate basin (the unbound state) and a sharp minimum (the bound state). We used the Chapman-Kolmogorov test to examine the MSM (Fig. S9C). While no clear violation of the expected transition probability is observed, the results of the test look atypical (and with the diagonal elements dominating fully) and likely arise from a poor convergence of the MSM due insufficient sampling of the slowest process. The latter hypothesis is supported by the observation that $t_1$ does not plateau within the range of considered lag-times (Fig. S9A).

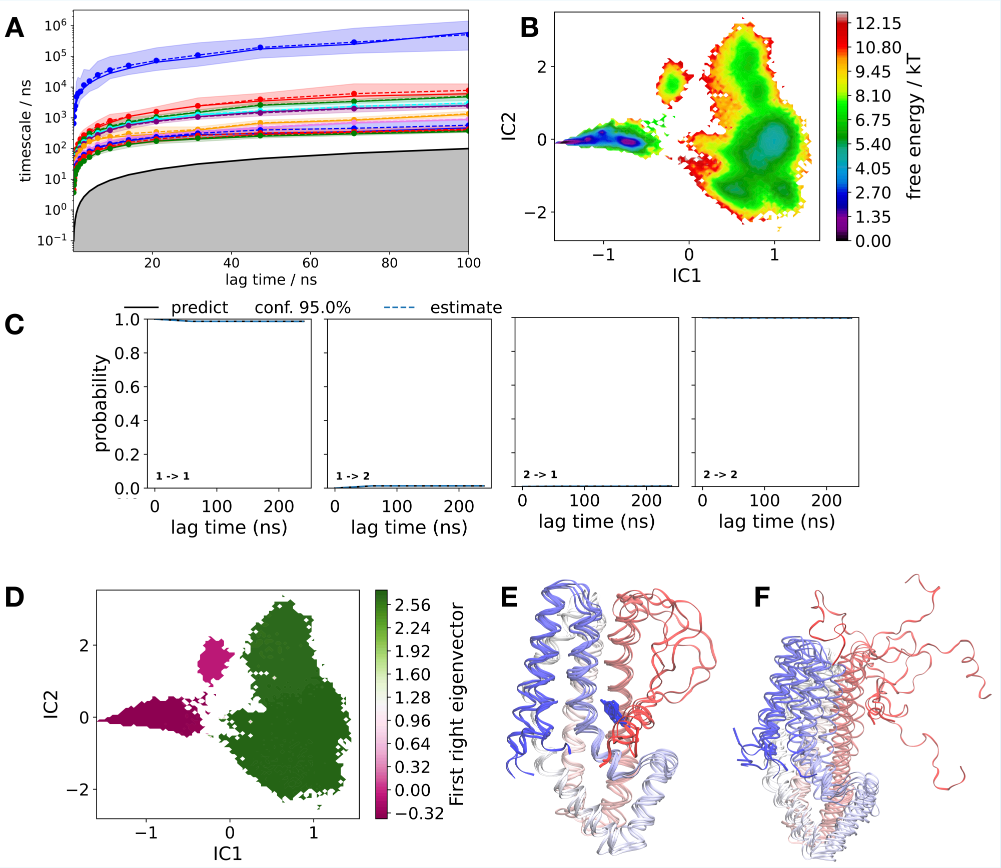


**Supporting Figure 9.** Constructing Markov State Models from the unbiased MD simulations. (A) The ten slowest implied timescales. The solid lines correspond to the implied timescales (and their 95% confidence intervals) of the maximum likelihood MSMs. The sample means are given by dashed lines and the grey area delimited by the solid black line represents timescales shorter than the lag-time. (B) The reweighted free energy obtained from the MSM by projecting onto the first two time-lagged independent components. (C) Results of the Chapman-Kolmogorov test on two metastable states. The black line represents the predicted transition probability $T_{ij}^{k}(\tau)$, where $T(\tau)$ is the transition probability matrix, $\tau$ is the lag-time, $k$ is an integer number and $i$ and $j$ are the two coarse-grained states, while the blue dashed line represents the estimated one, $T_{ij}(k\tau)$. (D) Projection onto the first two independent components of the first right eigenvector of the transition probability matrix. Renderings of the representative structures of the first (E) and second (F) cluster obtained by reducing the dimensionality of the MSM.

Analysis of the second right eigenvector of the transition probability matrix (Fig. S9D), which reports on the conformational transition that occurs at the $t_1$ timescale, clearly highlights the expected transition between the large degenerate basin and the sharper minimum (Fig. S9B); this is coherent with our expectations of the slow binding process. Finally, we used the robust Perron cluster cluster analysis (PCCA+) algorithm (***Röblitz and Weber, 2013***) to lump the MSM into a two-state model and sampled five representative structures from both clusters. These structures (Fig. S9E and S9F) represent, as expected, the bound and the unbound conformations of the Ac-helix. Further analyses of the MSM were prevented by the fact that it was impossible to build a meaningful Hidden Markov Model (HMM) (***Noé et al., 2013***) from our MSM because of the presence of two disconnected components in the transition probability matrix. This result was indeed expected due to, as we mentioned before, the absence of reversible transitions between states **2** and **3**. Building a more accurate MSM would thus require substantially more sampling focusing on this transition, which we did not attempt here due to the large computational requirements.

#### Populations of the bound and unbound states

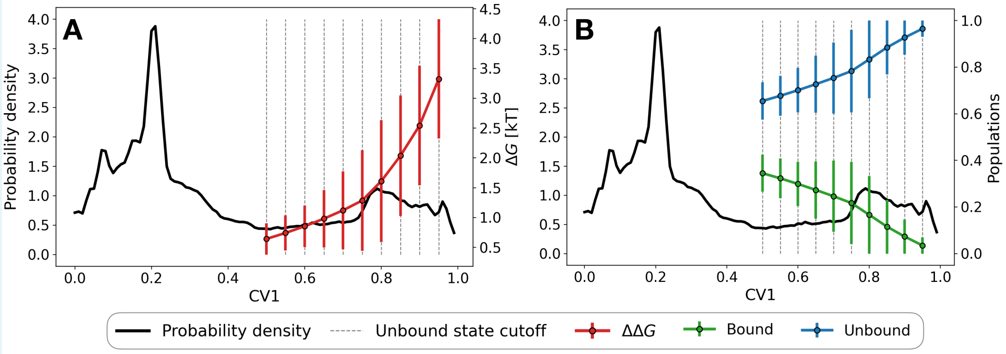


**Supporting Figure 10.** Block error analysis of the population of the unbound state. Probability density of CV1 in the PBMetaD simulation (black solid curve) and the different cutoffs used to determine the unbound state (dashed grey lines). For each cutoff, we report the value of (A) the $\Delta G_{\text{sim}}$ (red dot) and (B) the populations of the bound and unbound states (respectively, green and blue dots). Uncertainties were determined using block error analysis and, since the simulations were not fully converged, the reported values correspond to the error estimate using the two largest possible blocks (4 $\mu$s).

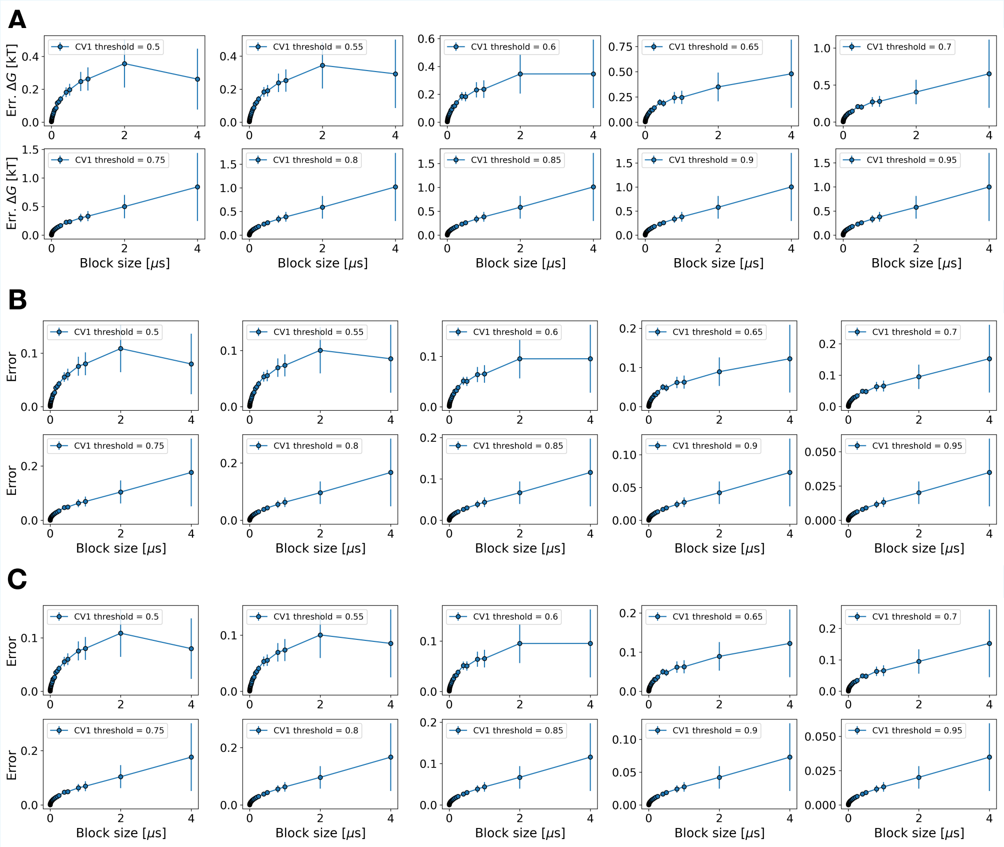


**Supporting Figure 11.** Error curves obtained by applying block error analysis to the (A) $\Delta G_{\text{sim}}$, (B) bound state population and (C) unbound state population obtained from different definitions of the unbound state. The upper threshold in CV1 used to determine the unbound state is reported in the figure labels.
